# Mechanistic characterization of tenuazonic acid-induced cellular stress responses in human esophageal KYSE-510 cells

**DOI:** 10.64898/2026.07.06.736731

**Authors:** Dino Grgic, Maximilian Jobst, Mariagiovanna Pais, Nazmi Waesoh, Sonja Hager, Giorgia Del Favero, Doris Marko

## Abstract

Tenuazonic acid (TeA) is an emerging *Alternaria* mycotoxin frequently detected in food and feed commodities, raising concerns about its toxicological relevance. Chronic oral exposure to TeA has been reported to induce dysplastic alterations in the esophageal mucosa of mice, while human biomonitoring data indicate an association between TeA exposure and esophageal cancer, although a causal relationship has not yet been established. At a mechanistic level, the effects of TeA in esophageal cells remain poorly characterized. Therefore, this study investigated the impact of TeA on cytotoxicity, oxidative stress, DNA damage, mitochondrial homeostasis, cell-cycle distribution and transcriptomic stress responses in human esophageal KYSE-510 cells. TeA induced a concentration-dependent reduction in metabolic activity and total protein content after 24 h exposure to 0.1-100 µM. Significant cytotoxicity was measured starting from 20 µM. At sub-cytotoxic concentrations, TeA triggered rapid ROS formation within 5-30 min exposure and induced formamidopyrimidine-DNA glycosylase (FPG) sensitive DNA damage after 1 h exposure (5-7.5 µM), indicating oxidative DNA lesions. In addition, TeA altered mitochondrial morphology after 4 h exposure at 7.5 µM, manifested by shrinkage of the mitochondrial network area and perinuclear redistribution, while mitochondrial respiration showed only a non-significant tendency towards reduced respiratory capacity. RNA sequencing after 6 h exposure to 10 µM TeA revealed oxidative stress-associated transcriptional changes, impaired antioxidant and stress-adaptive responses, and p53-associated stress signaling. Furthermore, TeA induced significant G₂/M phase accumulation after 24 h exposure to 1-10 µM.

**Highlights:**

- TeA triggers rapid oxidative stress in human esophageal KYSE-510 cells
- Sub-cytotoxic TeA induces FPG-sensitive oxidative DNA lesions
- TeA disrupts mitochondrial morphology before overt cytotoxicity occurs
- RNA-seq reveals impaired antioxidant defense and p53-linked stress signaling
- TeA promotes G₂/M phase accumulation and concentration-dependent cytotoxicity

**Graphical abstract:** 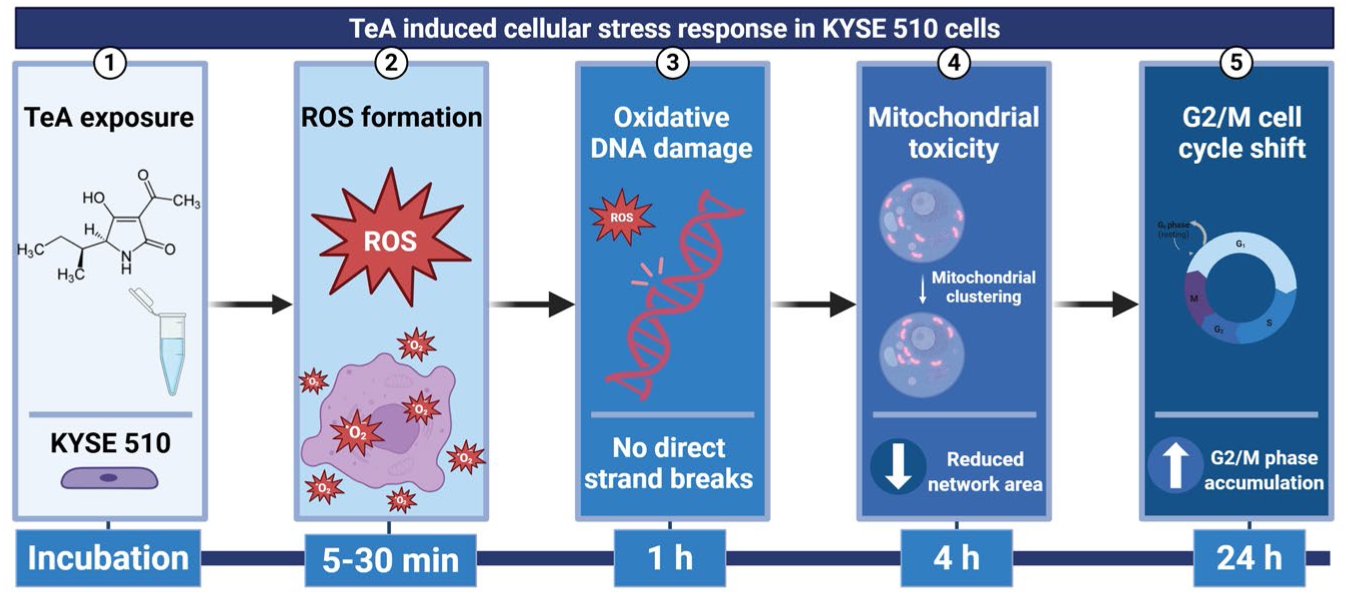

## Introduction

*Alternaria* species are ubiquitous molds that infect a broad range of crops. These organisms are capable of producing several secondary metabolites with the potential to induce adverse health effects. Consequently, these mycotoxins are relevant to the safety of food and feed. Tenuazonic acid (TeA), a 3-acyl tetramic acid derivative (Figure 1), is among the most relevant *Alternaria* toxins due to its frequent occurrence and comparatively high concentrations in contaminated commodities (Zhang, Liu et van der Fels-Klerx, 2025).

**Figure 1:**
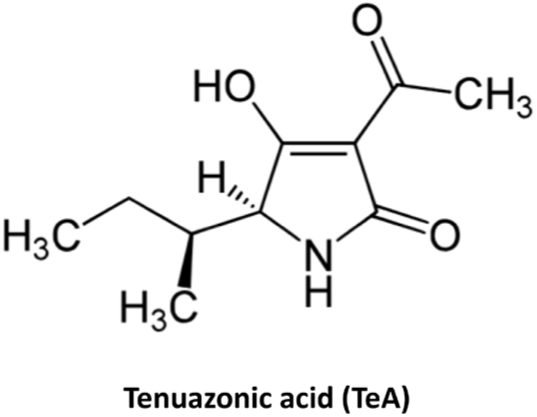
Structure of tenuazonic acid (TeA; CAS: 610-88-8)

Occurrence data demonstrate that TeA is frequently detected in food and feed, reaching comparatively high concentrations to other *Alternaria* mycotoxins. Relevant food commodities include cereals, tomato-based products, sunflower seeds, dried fruits, infant foods, herbs and spices (Asam et al. 2012; Asam et Rychlik 2013; Gotthardt et al. 2019; Louro et al. 2024).

Available absorption, distribution, metabolism and excretion (ADME) data indicate that TeA is readily absorbed and predominantly eliminated via the urine. In rats, the oral administration of *Alternaria* extracts resulted in the rapid recovery of TeA in the urine, supporting systemic uptake followed by renal excretion (Puntscher et al., 2019). Human biomonitoring studies identified urinary TeA as a relevant exposure marker for *Alternaria* toxins, as TeA was frequently detected in urine samples and showed high urinary recovery after the consumption of contaminated food (Hövelmann et al., 2016). Recent human toxicokinetic data further suggest possible phase I and phase II metabolism and rapid elimination of TeA after relevant systemic distribution in humans (Visintin et al., 2023).

A number of adverse effects have been associated with exposure to TeA. From a mechanistic perspective, early reports indicated potency to disrupt eukaryotic protein synthesis by impeding ribosomal function and hindering the release of nascent polypeptides from the ribosome (Carrasco et Vazquez 1972, 1973; Farber et al. 1971; Lieberman et al. 1970; Shigeura et Gordon 1963; Younger 1970). Furthermore, *in vivo* studies in mice have shown that intraperitoneal TeA exposure at 238 µg/kg for 8 weeks is associated with several physiological impairments, including oxidative stress, altered antioxidant enzyme activities, increased lipid peroxidation, changes in liver enzyme levels, and organ toxicity. In line with these systemic effects, additional adverse outcomes have been reported, including gastrointestinal bleeding, cardiovascular collapse, altered feed consumption, changes in organ weights, and decreased renal, splenic, cardiac, and brain weights (Kumari et Singh, 2021).

Despite this broad spectrum of reported toxicological effects, the available experimental and epidemiological data on the effects of TeA on the esophagus is limited. A potential toxic effect of TeA on esophageal tissue was indicated in a long-term mouse study. In this study, daily oral exposure to TeA at 25 mg/kg body weight for 10 months resulted in histopathological changes in the esophageal mucosa, including moderate to severe epithelial dysplasia, loss of cellular polarity, nuclear pleomorphism, and hyperchromasia (Yekeler et al., 2001). In a recent Ethiopian human biomonitoring case-control study, researchers indicated a positive correlation between TeA exposure and the risk of esophageal cancer. The study also revealed a higher frequency of TeA detection in plasma samples from esophageal cancer patients compared to controls (Mulisa et al., 2025). However, the currently available data do not allow a causal relationship to be established, as the human data are based on association and reflect co-exposure to multiple mycotoxins, while the animal data were obtained after high-dose chronic exposure. Thus, TeA might contribute to esophageal toxicity and potentially to esophageal carcinogenesis, but further mechanistic and prospective studies are required to clarify its specific role.

Therefore, the present study aimed to investigate the cellular effects of TeA in the human esophageal cell line KYSE-510, focusing on cytotoxicity, oxidative stress, DNA damage, mitochondrial alterations, cell-cycle perturbation and transcriptomic changes and its potential mechanistic contribution to chemical carcinogenesis.

## Methods

### Cell culture

KYSE-510 cells (human esophageal carcinoma cell model) were obtained from the German Collection of Microorganisms and Cell Cultures GmbH (DSMZ, Braunschweig, Germany). All cell culture reagents, including medium and supplements, were purchased from GIBCO Invitrogen (Karlsruhe, Germany). Cells were cultured in RPMI 1640 medium containing 10% fetal calf serum (FCS) and 1% penicillin/streptomycin, corresponding to final concentrations of 50 units/mL and 50 µg/mL, respectively. KYSE-510 cells were routinely passaged at approximately 80% confluency and used between passage number 5 and 17.

### Coupled CellTiter-Blue® and SRB cytotoxicity assay

KYSE-510 cells were seeded in 96-well plates at a density of 10,000 cells per well and incubated for 48 h to allow cell attachment. Subsequently, cells were treated with increasing concentrations of TeA (final concentration 0.1-100 µM; Santa Cruz Biotechnology, Dallas/TX, USA; OT: G0225; CAS: 610-88-8) or 0.1% DMSO (Carl Roth, Karlsruhe, Germany) as solvent control for 24 h. Stock solutions were prepared in DMSO at 1000-fold higher concentrations and diluted in cultivation medium immediately before treatment. All experiments were performed in at least three independent biological replicates, each measured in technical triplicates.

After 24 h of incubation, the medium was removed and 100 µL of CellTiter-Blue® (CTB) (Promega Corporation, Madison, WI, United States) incubation solution, prepared as a 1:10 dilution of CTB reagent in culture medium, was added to each well. Cells were incubated for 60 min with the CTB solution, and fluorescence was measured using an excitation wavelength of 560 nm and an emission wavelength of 590 nm with a gain of 80 using the Cytation 3 Cell Imaging Multi-Mode Reader from BioTek® (Winooski/VT, USA). Results were normalized to the solvent control and expressed as percentage of control.

Following CTB measurement, cells were fixed with 10 µL of 50% (w/v) trichloroacetic acid in distilled water for 1 h at 4 °C. Plates were washed four times with tap water and left to dry overnight in the dark. Subsequently, 50 µL of SRB staining solution, prepared by dissolving 4 g SRB reagent in 1 L distilled water containing 1% acetic acid, was added to each well. After staining for 1 h at room temperature in the dark, the staining solution was removed, and plates were washed twice with tap water followed by two washing steps with 1% acetic acid. Plates were again dried overnight in the dark, and the protein-bound dye was dissolved under alkaline conditions by adding 100 µL Tris base solution (0.30 g tris(hydroxymethyl)aminomethane in 250 mL distilled water) and shaking for 5 min. Absorbance was measured at 570 nm using the Cytation 3 Cell Imaging Multi-Mode Reader from BioTek® (Winooski/VT, USA). Final results were normalized to the solvent control and expressed as percentage of control.

### Dichlorofluorescein diacetate (DCF-DA) assay

The formation of intracellular reactive oxygen species (ROS), as marker for oxidative stress, was assessed using the DCF-DA assay according to Keston and Brandt (1965). The assay relies on the uptake of the non-fluorescent dye dichlorodihydrofluorescein diacetate into the cells, followed by intracellular de-esterification to dichlorodihydrofluorescein. In the presence of ROS, this compound is subsequently oxidized to the fluorescent product 2′,7′-dichlorofluorescein. KYSE-510 cells (10,000 cells/well) were seeded in black 96-well plates with clear bottoms and cultivated for 48 h before treatment. All subsequent steps were performed under light-protected conditions using red light to avoid photooxidation.

Cells were washed once with phosphate-buffered saline (PBS) and incubated with 100 µL DCF-DA solution (50 µM in colorless serum-free RPMI medium) for 15 min at 37 °C. After careful aspiration and two washing steps with PBS, background fluorescence was measured. Cells were then treated in colorless serum-free medium with the indicated TeA concentrations or hydrogen peroxide (H_2_O_2_) as positive control. Fluorescence was recorded at defined time points between 5 and 30 min using the Cytation 3 Cell Imaging Multi-Mode Reader from BioTek® (Winooski/VT, USA) at excitation/emission wavelengths of 485/528 nm. Fluorescence values were corrected for detector gain based on the positive control. Cell integrity was visually checked after the measurement to exclude cell loss during washing. Oxidative stress was calculated as relative fluorescence and expressed as percentage of the solvent control according to the following equation: oxidative stress (%) = 100 × emission of treated sample / emission of control.

### Comet assay

DNA strand breaks and oxidative DNA damage were assessed using the alkaline comet assay, performed according to Tice et al. (2000). For this purpose, 150,000 KYSE-510 cells were seeded in 3.5 cm petri dishes and cultivated for 48 h to allow attachment. Cells were then exposed to the respective test compounds or controls for 1 h. After incubation, cells were washed with PBS, detached by trypsinization and counted. Cell viability was determined by trypan blue exclusion and only samples with a viability above 80% were further processed.

For each condition, two slides were prepared. Therefore, 30,000 cells were embedded in agarose in duplicate on each slide. To remove cellular components and allow DNA unwinding, slides were incubated overnight at 4 °C in lysis buffer containing 1% lauryl sarcosinate, 1% Triton X-100 and 10% DMSO. On the following day, slides were washed three times with formamidopyrimidine-DNA glycosylase (FPG) buffer consisting of 40 mM HEPES, 100 mM KCl, 500 µM EDTA and 20 mg/mL bovine serum albumin at pH 8.0. To additionally detect FPG sensitive sides, slides were incubated either with FPG buffer alone or with FPG enzyme solution (50 µL of 4 U/mL in FPG buffer per slide) for 30 min at 37 C.

Subsequently, slides were incubated in alkaline electrophoresis buffer at pH 13 for 20 min to allow DNA unwinding, followed by electrophoresis for 20 min at 25 V, corresponding to 0.028 V/cm², and 300 ± 3 mA. After electrophoresis, slides were neutralized and stained with ethidium bromide at a concentration of 0.2 mg/mL. DNA migration was analyzed using a Zeiss Axioskop fluorescence microscope with an excitation wavelength of 546 ± 1 nm and an emission wavelength of 590 nm. Tail intensity was quantified with Comet Assay IV software (Perceptive Instruments). In total, 50 nuclei per pad, corresponding to 100 nuclei per slide, were evaluated and the mean tail intensity was used as a marker of DNA fragmentation.

### Live cell imaging

KYSE-510 cells were seeded in µ-Slide 8 Well ibiTreat chambers (REF: 80.826, ibidi GmbH, Gräfelfing, Germany) at a density of 10,000 cells per well and allowed to adhere for 48 h. Cells were treated with TeA (1-7.5 µM) or 0.1% DMSO for 4 h. After incubation, cells were washed with warm normal external solution (NES) (Del Favero et al., 2012) and stained for 15 min with MitoTracker™ (Invitrogen; REF: M7514) at a dilution of 1:1000. Cells were washed once and imaged in NES. Live cell imaging was carried out using the Lionheart FX Automated Microscope from BioTek® (Vermont, USA; RRID: SCR_019744). Images were acquired with the GEN5 Microplate Reader and Imager Software version 3.05 from BioTek® (Vermont, USA; RRID: SCR_017317). Mitochondrial area was quantified using ImageJ 1.54g.

### Seahorse assay

Mitochondrial respiration was assessed using the Seahorse XF assay according to the manufacturer’s instructions and previously published protocols (Jobst et al., 2025; Mayberry et al., 2024). Briefly, KYSE-510 cells were seeded in Seahorse XF 24 well cell culture plates at a density of 7500 cells per well in RPMI cultivation medium supplemented with 10% FBS and 1% P/S, resulting in a basal oxygen consumption rate of approximately 30 pmol/min. Wells containing medium only were included as blanks. After seeding, cells were kept for 1 h at room temperature to allow proper attachment and were then incubated for 24 h at 37 °C and 5% CO_2_.

On the same day, the sensor cartridge was hydrated with 1 mL XF calibrant per well and incubated for 24 h at 37 °C in a CO_2_-free incubator. On the following day, cells were treated with TeA (1-7.5 µM) or 0.1% DMSO for 4 h. Seahorse assay medium was freshly prepared by supplementing 19.4 mL Seahorse XF DMEM with 0.2 mL Seahorse XF glucose (final concentration: 25 mM), 0.2 mL Seahorse XF pyruvate (final concentration: 1 mM) and 0.2 mL Seahorse XF glutamine (final concentration: 2 mM). After treatment, medium was removed and 500 µL Seahorse assay medium was added to each well, followed by incubation for 1 h at 37 °C in a CO_2_-free incubator.

In parallel, the sensor cartridge was loaded with the respective mitochondrial modulators in the following order and final concentrations: 3 µM oligomycin, 3 µM carbonyl cyanide m-chlorophenyl hydrazone (CCCP), 5 µM antimycin A and 5 µM rotenone. At the beginning of each measurement, three mixing and measuring cycles were performed to determine basal respiration. Subsequently, the mitochondrial modulators were injected sequentially, and the mixing and measuring cycles were repeated after each injection. Oxygen consumption rate (OCR) was measured using a Seahorse XFe24 analyzer (Seahorse Bioscience, USA). OCR data were analyzed using OriginPro software.

### Cell cycle analysis

KYSE-510 cells were seeded in 6-well plates at a density of 150,000 cells per well and cultivated for 24 h. Cells were then treated for another 24 h with TeA at final concentrations of 1-10 µM, 0.1% DMSO or 5 µM oxaliplatin (Ox) as positive control. After treatment, cells were harvested by trypsinization, washed with PBS and fixed in 70% ice-cold ethanol at -20 °C overnight. The following day, suspensions were centrifuged at 1000 × g for 10 min, the supernatant was discarded, and the cell pellet was washed twice with PBS and centrifuged again to remove residual ethanol.

Cell pellets were then resuspended and stained in PBS containing ribonuclease (RNase; 330 µg/mL RNase A and H) and propidium iodide (PI; 10 µg/mL) for 30 min at room temperature. Samples were analyzed using a Guava easyCyte flow cytometer. Allocation of cells to the respective cell cycle phases and quantitative analysis were performed using Floreada.io software.

### RNA sequencing

KYSE-510 cells were seeded at a density of 150,000 cells per well (12-well plates) and cultured for 48 h. Incubation solutions containing 0.1% DMSO (solvent controls) and the respective test compounds or compound mixtures were subsequently prepared and applied to the cells. Following 6 h of exposure, cells were washed and total RNA was isolated using the RNeasy Mini Kit (Qiagen) according to the manufacturer’s instructions. RNA quality control, library preparation, and sequencing procedures were performed by Novogene (Planegg, Germany). RNA quality, purity, and integrity were assessed using Qubit, NanoDrop, and Bioanalyzer/TapeStation systems. Subsequently, cDNA libraries were generated using automated workstation platforms and sequenced on Illumina NovaSeq platforms.

All preprocessing steps were performed on a high-performance computing cluster. Raw paired-end reads were quality-checked using FastQC (Andrews, 2010) and MultiQC (Ewels et al., 2016), trimmed with Cutadapt (Saeidipour et Bakhshi 2013), and quality control was repeated after trimming. Trimmed reads were aligned to the human reference genome GRCh38, Ensembl release 115, using STAR (v2.7.3a) (Dobin et al., 2013). Gene-level read counts were generated with HTSeq (Anders et al., 2015) in strand-specific mode using the Ensembl GRCh38.115 annotation, and merged into gene-by-sample count matrices separately.

Downstream analyses were performed in R (v4.3+)(Team, 2014). Differential expression analysis was performed using DESeq2 (v1.40+) (Love et al., 2014), with treatment concentration as the design factor and DMSO as the reference. Library size differences were corrected using the median-of-ratios normalisation method, statistical testing was performed using the Wald test, and p-values were adjusted using the Benjamini-Hochberg procedure. For each contrast, full results tables were exported and used for downstream functional enrichment analysis.

Sample-level transcriptomic variation was assessed by PCA based on VST-normalised expression data. MA plots were generated with DESeq2, and volcano plots were created using ggplot2 (Wickham, 2016). The top genes by adjusted p-value were labelled using ggrepel (Slowikowski et al., 2024).

Functional enrichment analysis was performed using clusterProfiler (v4.x) (Wu et al., 2021; Yu et al., 2012) and ReactomePA (Yu et He 2016), with gene annotation from org.Hs.eg.db (Carlson et al., 2019). Genes were selected based on adjusted p-value and converted from Ensembl IDs to Entrez IDs using the bitr function. Gene ontology (GO) enrichment analysis was performed for Biological Process (BP). Enrichment results were ordered by significance.

### Statistics

The CTB, SRB, DCF-DA, and Seahorse assay were performed in at least biological triplicates (n ≥ 3) with technical triplicates. The comet assay and live cell imaging were performed in biological triplicates with technical duplicates. For live cell imaging, a total of 24 images were acquired for each condition, corresponding to 8 randomly chosen optical fields for each cell preparation (biological replicate). Four cells per image were used for quantification, resulting in 96 cells per condition. Cell cycle analysis and RNA sequencing were performed in three biological replicates without technical replicates. Statistical analysis and data visualization were performed using OriginPro® 2026. Significance levels were defined as 5%, 1% and 0.1%, respectively (* p < 0.05; ** p < 0.01; *** p < 0.001). Significant differences were calculated by one-way analysis of variance (ANOVA) followed by Bonferroni post hoc test.

## Results

### Cytotoxicity

Treatment with TeA led to a concentration-dependent decrease in metabolic activity, as measured by the CTB assay (Figure 2A). Concentrations as low as 10 µM did not result in a statistically significant change in metabolic activity. Starting at 20 µM, metabolic activity exhibited a significant decrease compared to 0.1% DMSO, reaching 67 ± 15% and 62 ± 12% at 20 and 25 µM, respectively. A further decline in metabolic activity was observed at higher concentrations of 75 µM and 100 µM, reaching 47 ± 10% and 44 ± 12%, respectively. The Triton^TM^ X-100 (Triton-X) control, utilized to induce cell death, confirmed loss of metabolic activity to 15 ± 12%.

**Figure 2:**
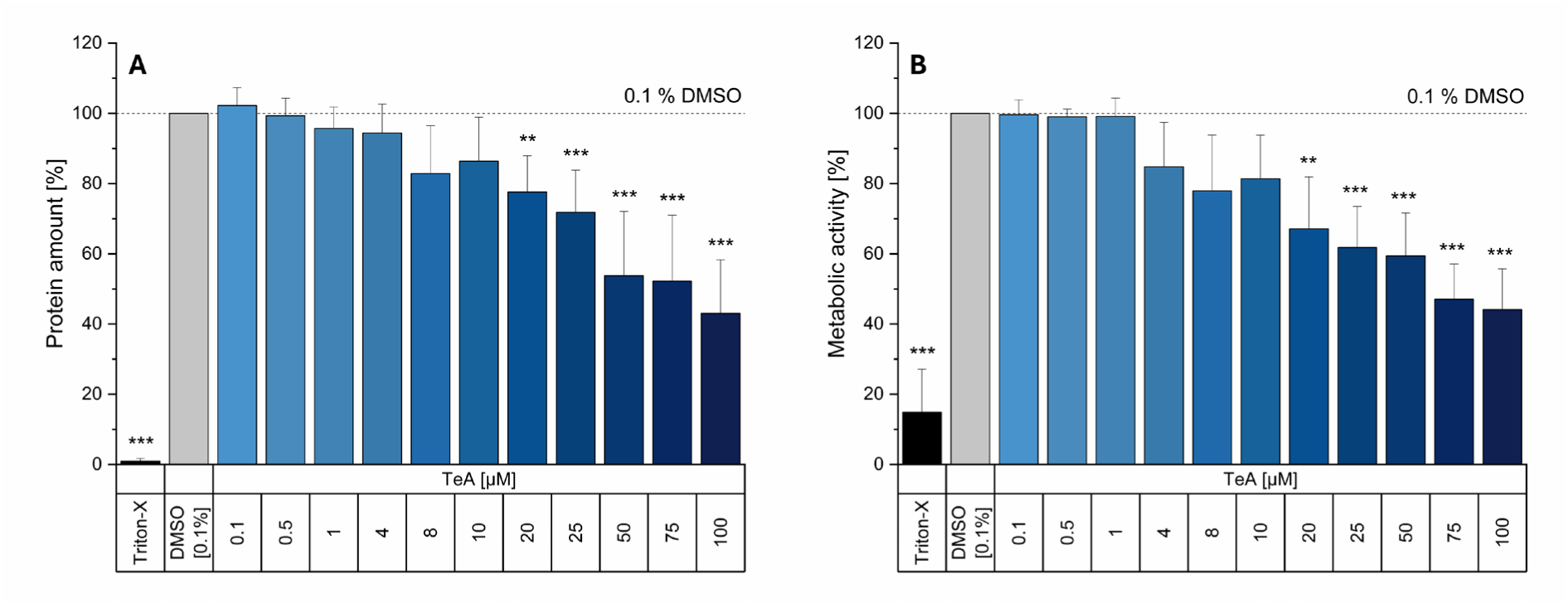
Effects of tenuazonic acid (TeA) on the cytotoxicity of KYSE-510 cells. Impact on the cell viability [%] measured by the CellTiter Blue (CTB) assay (A) and on the cell protein amount [%] measured by the sulforhodamine B (SRB) assay (B) of increasing concentrations of TeA (0.1–100 µM), 0.1% DMSO and 0.01% Triton-X after 24 h incubation in KYSE-510 cells. Values were normalized to solvent control (0.1% DMSO) and are depicted as mean ± SD of at least 3 independent biological replicates, each measured in technical triplicates. Significant differences of effects between the solvent control and TeA were calculated by one-way ANOVA followed by Bonferroni post hoc test and are indicated with **p < 0.01 and *** p < 0.001.

Similarly, total cellular protein content measured by the SRB assay showed a concentration dependent reduction following TeA exposure (Figure 2B). Protein levels remained largely unaffected at concentrations up to 10 µM, whereas a gradual decrease became evident from 10 µM onwards. A significant decrease in protein content was observed at 20 µM, with a reduction to 77 ± 10%. With increasing concentrations of 50 and 75 µM TeA, the protein amount further declined to 54 ± 18% and 52 ± 19%, respectively. The decline was most pronounced at the highest applied concentration of 100 µM, resulting in a reduction of 43 ± 15% compared to the solvent control (0.1% DMSO).

### ROS formation

Intracellular ROS formation was assessed using the DCF-DA assay following TeA exposure at different concentrations and time points (Figure 3). The positive control H₂O₂ induced a pronounced and significant increase in fluorescence to 177 ± 8%, 230 ± 29% and 252 ± 23% at the time points 5, 15 and 30, respectively.

**Figure 3:**
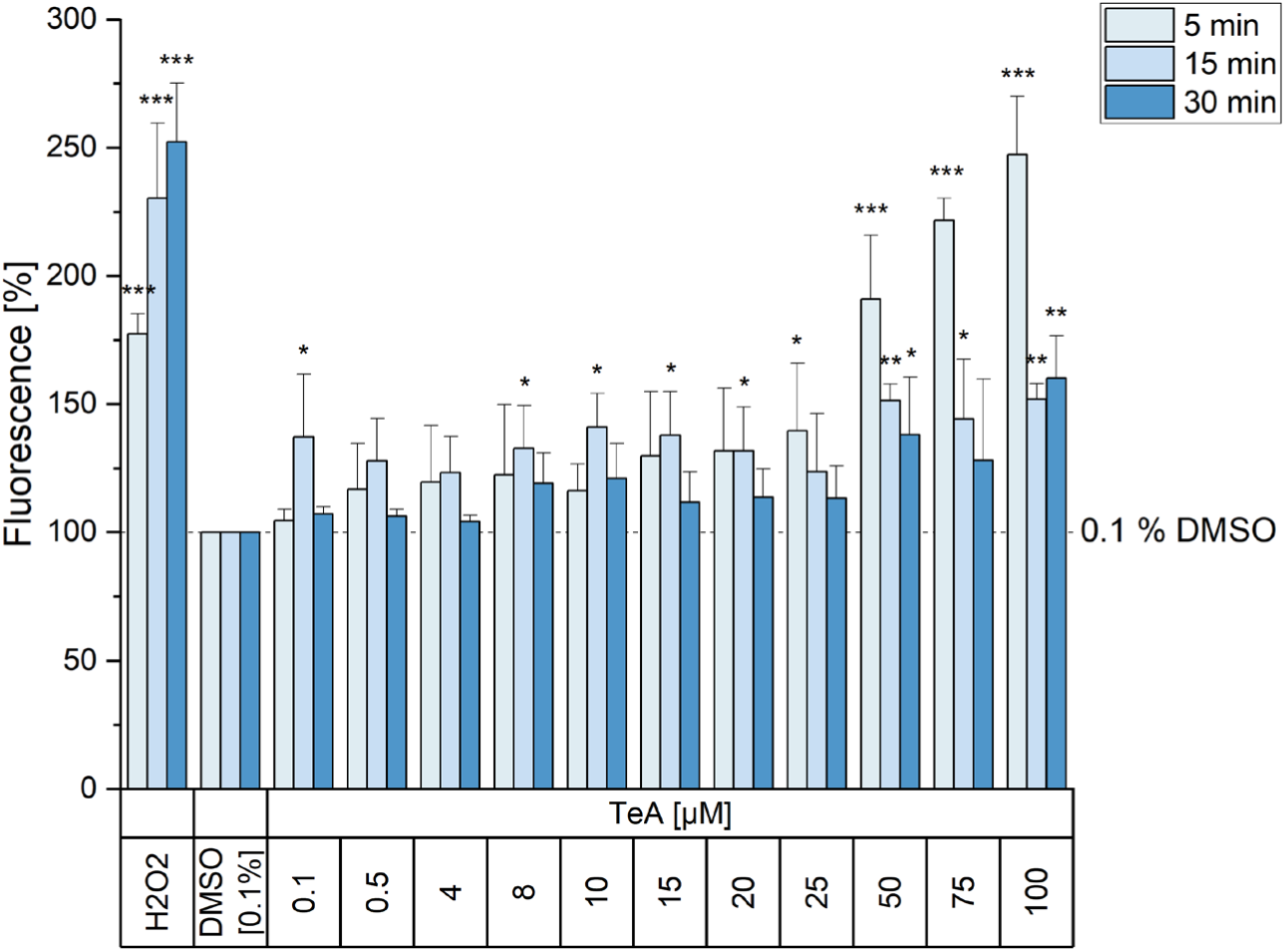
Results of the Dichlorofluorescein diacetate (DCF-DA) assay in KYSE-510 cells incubated for 5, 15 and 30 minutes in serum-free colorless medium with increasing concentrations of tenuazonic acid (TeA; 0.1–100 µM) and the positive control hydrogen peroxide (H_2_O_2_; 1 mM). Values were normalized to solvent control (0.1% DMSO) and are depicted as mean ± SD of at least 3 independent biological replicates, each measured in technical triplicates. Significant differences of effects between the solvent control and TeA were calculated by one-way ANOVA followed by Bonferroni post hoc test and significances are indicated with * p < 0.05, ** p < 0.01 and *** p < 0.001.

After 5 minutes incubation, low to intermediate TeA concentrations (0.1-20 µM) caused a slight concentration-dependent but non-significant increase in fluorescence intensity compared to the solvent control. A marked and significant elevation of ROS formation was observed at higher concentrations during the initial measurement (25-100 µM, 5 min). At 15 min, only low to intermediate TeA concentrations resulted in an additional increase in ROS formation compared to the previous time point at 5 min. Compared with the solvent control (0.1% DMSO), ROS levels were statistically significantly increased at 0.1 µM and at 8-20 µM TeA. Starting at 25 µM (15 min), a decline in ROS formation was observed in comparison with the initial time point (5 min). At 30 minutes, a decrease was evident for all concentrations when compared with the earlier time points (5 and 15 min).

### Comet assay

The alkaline comet assay was used to assess DNA damage in KYSE-510 cells after 1 h of TeA exposure. DNA damage was quantified as tail intensity [%], reflecting the amount of DNA migrating from the nucleus during electrophoresis. To further evaluate oxidative DNA damage, the assay was performed in the absence and presence of formamidopyrimidine DNA glycosylase (FPG), which converts oxidized purines into additional DNA strand breaks detectable by the comet assay. The UV-light treated positive control showed a strong and significant increase in tail intensity both in the absence and presence of FPG, confirming the sensitivity of the assay (Figure 4). After 1 h of incubation, TeA induced DNA damage in KYSE-510 cells mainly after FPG treatment. In the absence of FPG, tail intensity remained low across all tested TeA concentrations and was comparable to the solvent control. No concentration-dependent increase in DNA strand breaks was observed under these conditions. In contrast, FPG treatment resulted in a clear increase in tail intensity, especially at 5 and 7.5 µM TeA to 13 ± 3 and 16 ± 3%, respectively. At these concentrations, tail intensity was significantly elevated compared to the corresponding control, indicating the formation of FPG-sensitive DNA lesions.

**Figure 4:**
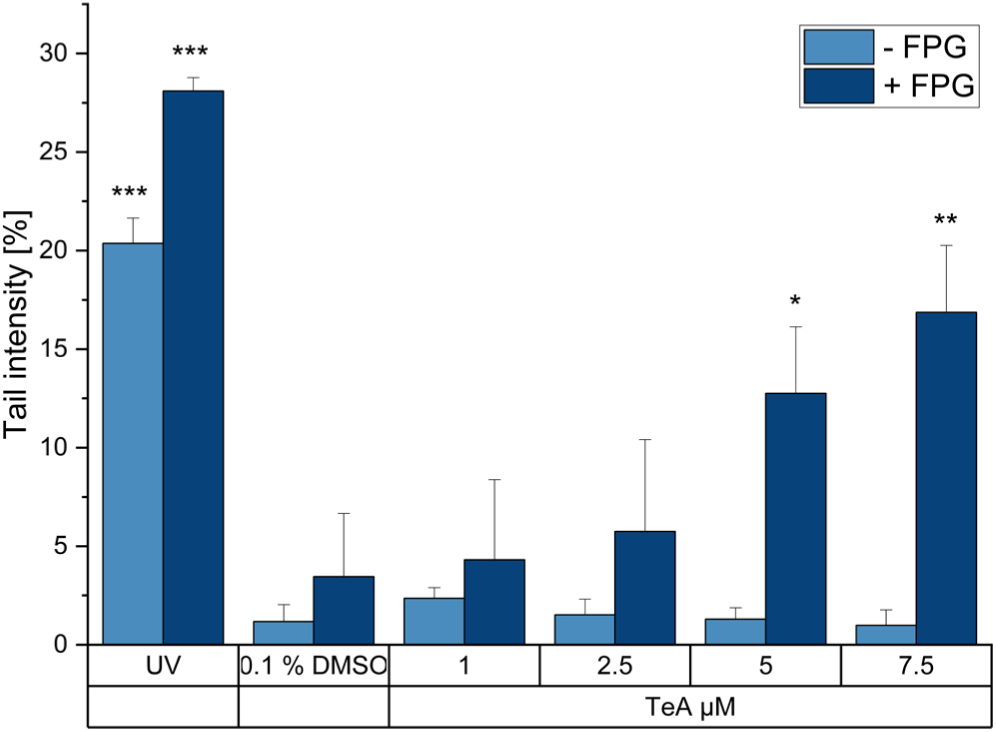
Results of the alkaline comet assay in KYSE-510 cells incubated for 1 h with increasing concentrations of tenuazonic acid (TeA) and the positive control UV, with and without formamidopyrimidine DNA glycosylase (FPG) treatment. DNA damage is expressed as tail intensity [%]. Values are depicted as mean ± SD of 3 independent biological replicates. Significant differences of effects between the solvent control (0.1% DMSO) and TeA or UV were calculated by one-way ANOVA followed by Bonferroni post hoc test and significances are indicated with * p < 0.05, ** p < 0.01 and *** p < 0.001.

### Mitochondrial morphology

Mitochondrial morphology was assessed by quantifying the area of the MitoTracker fluorescence signal following TeA exposure (Figure 5). Compared to the solvent control, low concentrations of TeA (1-5 µM) did not result in significant alterations of the mitochondrial intracellular distribution compared to 0.1% DMSO. In contrast, treatment with 7.5 µM TeA led to a significant reduction in the mitochondrial signal area, indicating a decrease in mitochondrial network size or density after 4 h of incubation (Figure 5 A and B). Mitochondrial shift toward the nuclei was visible, and the quantification indicated a significant decrease of the mitochondrial network area at 7.5 µM TeA, while the cell size and the signal intensity of the network remained constant (Figure 5C & D).

**Figure 5:**
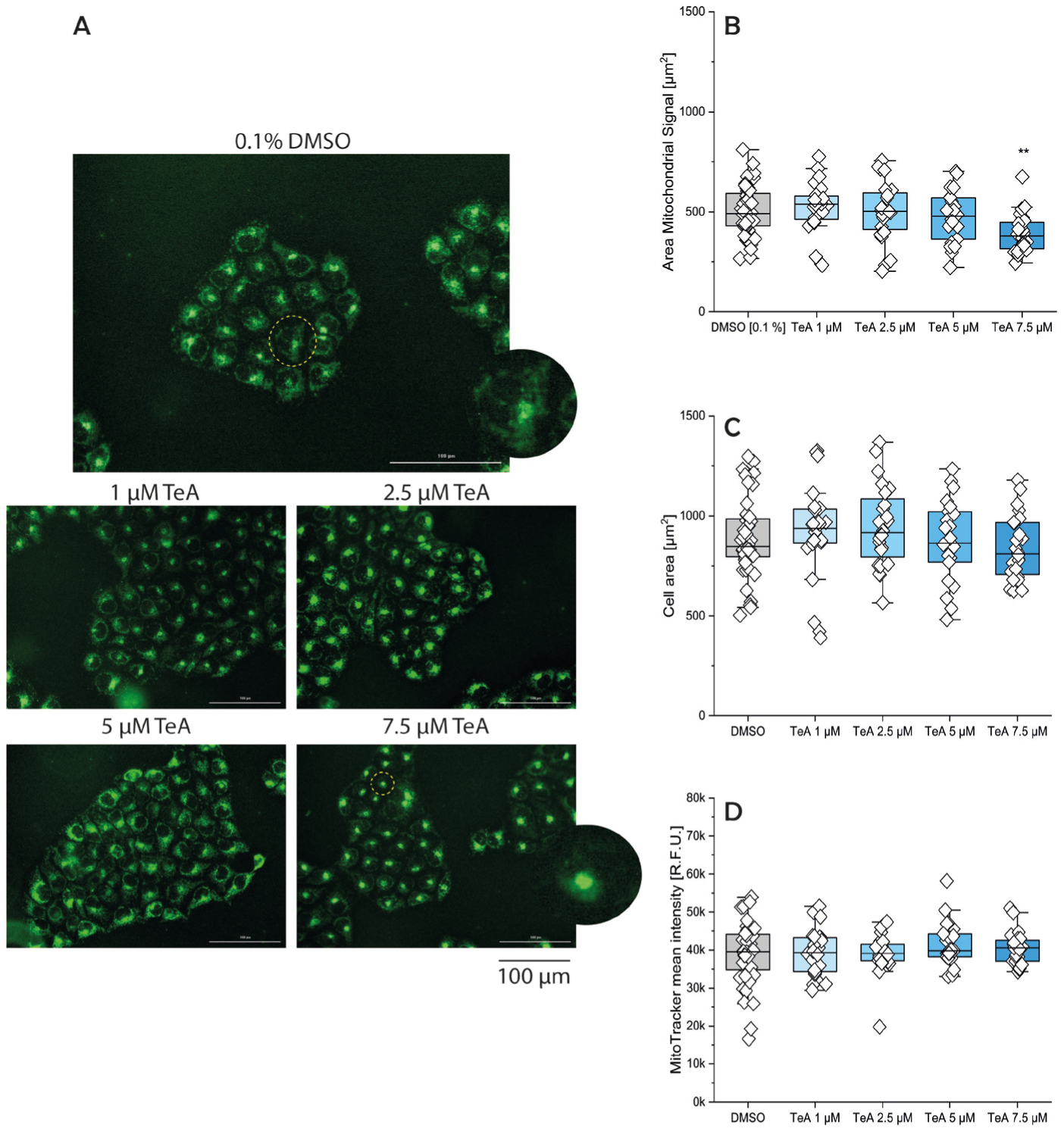
Representative images of mitochondrial network (green) in KYSE-510 cells incubated at 37°C with 0.1% DMSO and tenuazonic acid (TeA; 1-7.5 µM) for 4 h, scale bar 100 μm (A). Mitochondrial area was quantified as the average mitochondrial signal per optical field (B). Cell area was quantified as the average cell area per optical field (C). MitoTracker signal intensity was quantified as the average fluorescence intensity of the MitoTracker signal per optical field (D). Results are shown as boxplots, whiskers represent SD, and boxes represent the range from 25 to 75 percentage. Values are depicted as mean ± SD of 3 independent biological replicates, each measured in technical duplicates. For each technical replicate, four images were acquired, and four cells per image were used for evaluation. Significant differences of effects between 0.1% DMSO and TeA were calculated by one-way ANOVA followed by Bonferroni post hoc test and significances are indicated with **p < 0.01.

### Oxygen consumption rate (OCR)

Mitochondrial respiration was analyzed by measuring the OCR using the Seahorse XF assay following TeA exposure (Figure 6). Under basal conditions, control cells exhibited stable OCR values of approximately 31-33 pmol/min. Treatment with TeA in concentrations between 1-5 µM did not have an impact on the OCR compared to 0.1% DMSO. A slight decrease in OCR was observed after incubation with 7.5 µM TeA. However, this reduction was not statistically significant when compared to solvent control.

**Figure 6:**
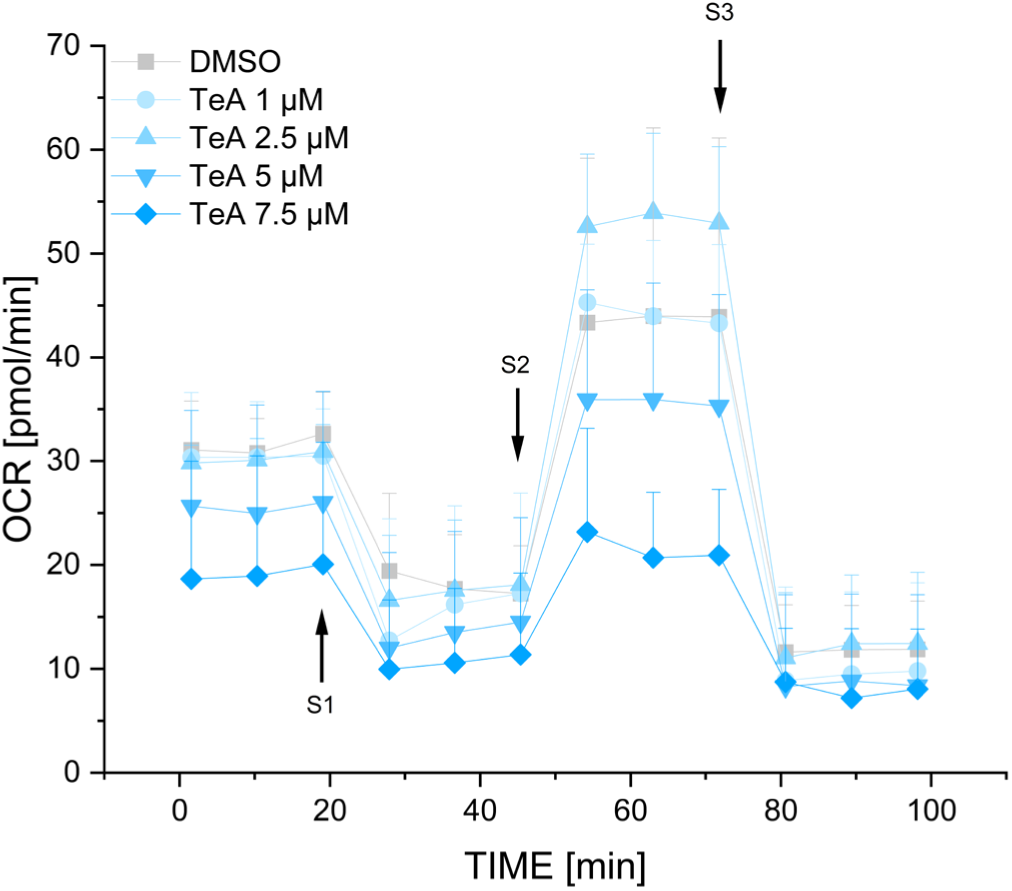
Seahorse assay in KYSE-510 cells pretreated for 4 h with 0.1% DMSO or increasing concentrations of tenuazonic acid (TeA; 1-7.5 µM). The time-dependent changes in oxygen consumption rate (OCR) are shown over the course of the measurement. After determination of basal respiration, 3 µM oligomycin was injected as first stimulus (S1), followed by 3 µM carbonyl cyanide m-chlorophenyl hydrazone (CCCP) as second stimulus (S2), and a combination of 0.5 µM antimycin A and rotenone as third stimulus (S3). Values are depicted as mean ± SD of at least 3 independent biological replicates, each measured in technical triplicates.

Following the administration of oligomycin (S1), which inhibits ATP synthase and thereby allows the assessment of ATP-linked respiration, OCR decreased markedly under all treatment conditions. Compared with the solvent control (0.1% DMSO), TeA dampened OCR in a concentration-dependent manner, although this effect was not statistically significant. Subsequent addition of the uncoupler of oxidative phosphorylation CCCP (S2) induced a marked increase in OCR in control cells, representing maximal respiratory capacity. While cells treated with low TeA concentrations (1-5 µM) still showed a comparable increase in OCR compared to 0.1% DMSO following CCCP injection, the maximal respiration was diminished at a TeA concentration of 7.5 µM. However, this reduction was not statistically significant. Finally, injection of rotenone and antimycin A (S3), inhibitors of mitochondrial complexes I and III, led to a strong reduction of OCR in all conditions, confirming effective inhibition of mitochondrial respiration and revealing non-mitochondrial oxygen consumption. Residual OCR values were comparable across treatments, indicating that the observed differences in earlier phases were attributable to mitochondrial activity.

### Cell cycle analysis

Cell cycle distribution was subsequently analyzed by flow cytometry after 24 h of TeA exposure and compared with the solvent control (Figure 7). In control conditions (0.1% DMSO), cell distribution was predominantly observed between the G_0_/G_1_ phase (52 ± 6%) and S phases (36 ± 14%), with a comparatively smaller proportion of cells in the G_2_/M phase (8 ± 3%).

**Figure 7:**
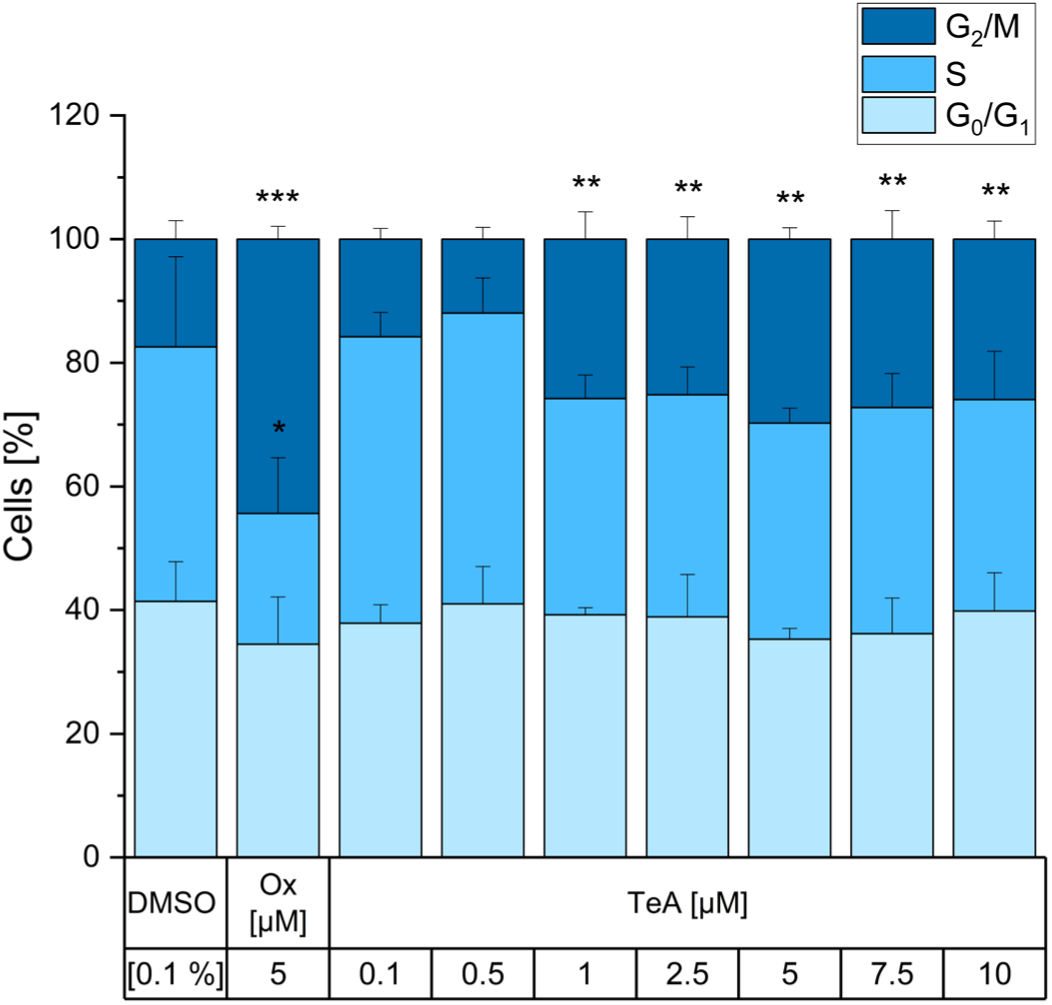
Cell cycle changes of KYSE-510 cells upon treatment with 0.1% DMSO, 5 µM Oxaliplatin (Ox) as a control to induce G_2_/M arrest and tenuazonic acid (TeA; 1-7.5 µM). Values are depicted as mean ± SD of at least 3 independent biological replicates. Significance was calculated using 2-way ANOVA followed by Bonferroni post hoc test (* p < 0.05, ** p < 0.01, *** p < 0.001).

Treatment with 5 µM oxaliplatin (Ox), applied to induce G_2_/M arrest, resulted in a substantial shift in cell cycle distribution. This led to a significant accumulation of cells in the G_2_/M phase (44 ± 2%), accompanied by a concomitant decrease of cells in the G_0_/G_1_ (34 ± 8%) and S phase (21 ± 9%). Exposure to TeA in concentrations ranging from 1 to 10 µM resulted in alterations in cell distribution compared with 0.1% DMSO. The administration of TeA did not result in statistically significant alterations in the G_0_/G_1_ and S phase compared with the solvent control. In contrast, TeA at concentrations of 1 to 10 µM resulted in a significant increase in G_2_/M phase (25 – 29%).

### RNA sequencing: GO enrichment analysis for BP

For the interpretation of the RNA-seq data, GO-BP terms were selected based on their relevance to the experimentally investigated endpoints, rather than solely according to their overall enrichment significance.

#### Oxidative stress-related pathways

To investigate transcriptomic changes induced by TeA, KYSE-510 cells were exposed to 1 and 10 µM TeA or the solvent control (0.1% DMSO) for 6 h, followed by total RNA isolation and RNA sequencing. After preprocessing, alignment to the human reference genome, and differential gene expression analysis, GO-BP enrichment analysis was performed to identify significantly enriched biological processes among the differentially expressed genes. The GO-BP term *response to oxidative stress* was significantly enriched in KYSE-510 cells following exposure to 10 µM TeA compared with DMSO controls (p.adjust = 8.17 × 10⁻⁴) (Table 1). The corresponding enrichment plot (Figure 8) illustrates the overall enrichment of GO-BP term *response to oxidative stress* (GO:0006979) within the ranked gene list.

**Figure 8:**
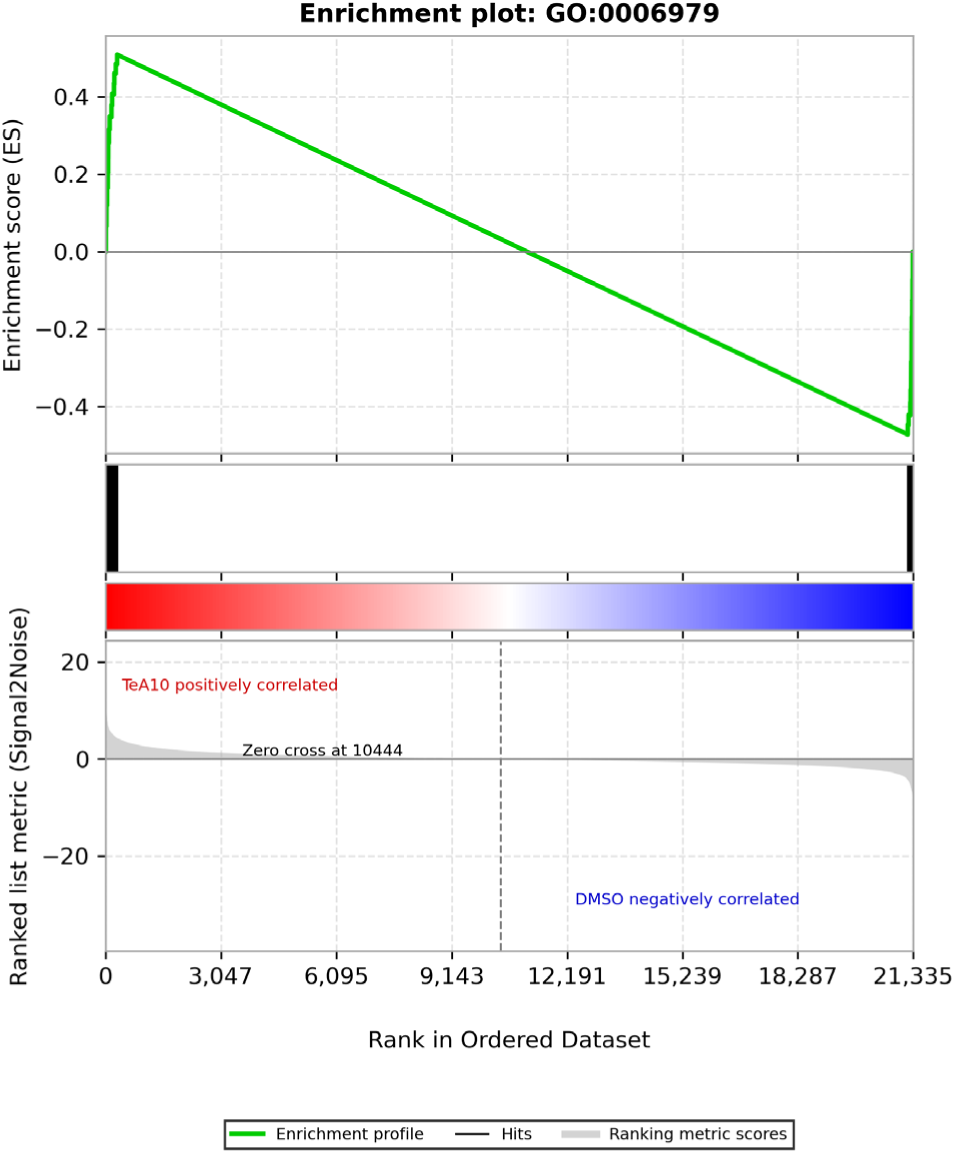
Gene set enrichment plot of the GO-BP term response to oxidative stress (GO:0006979) in KYSE-510 cells after exposure to 10 µM tenuazonic acid (TeA) for 6 h compared with 0.1% DMSO control. Genes are ranked according to the signal-to-noise metric, with genes positively correlated with 10 µM TeA located on the left and genes negatively correlated with DMSO located on the right. Black vertical lines indicate the position of genes belonging to the GO:0006979 gene set within the ranked dataset, and the green line represents the running enrichment score. The plot illustrates the enrichment of oxidative stress response-associated genes toward the 10 µM TeA-correlated side of the ranked gene list.

**Table 1:**
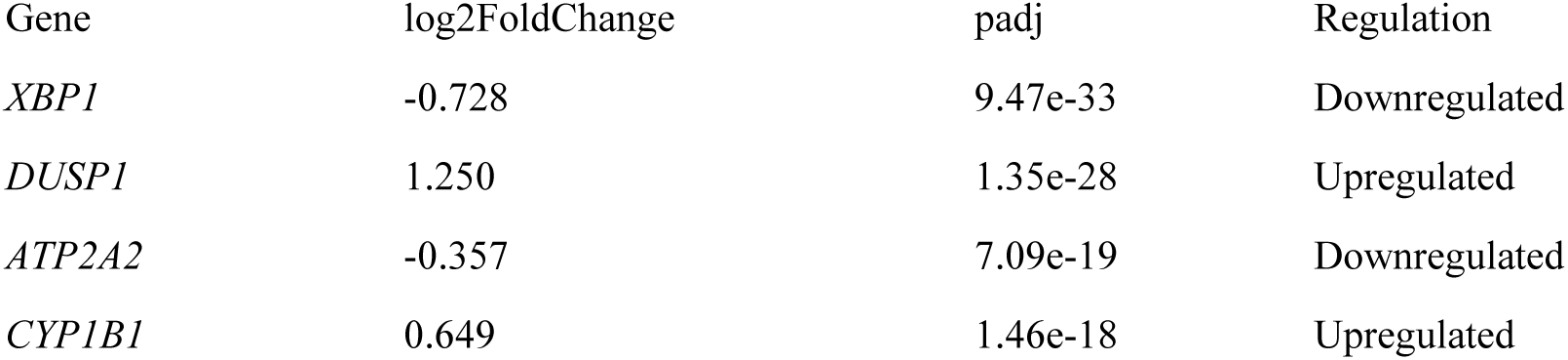
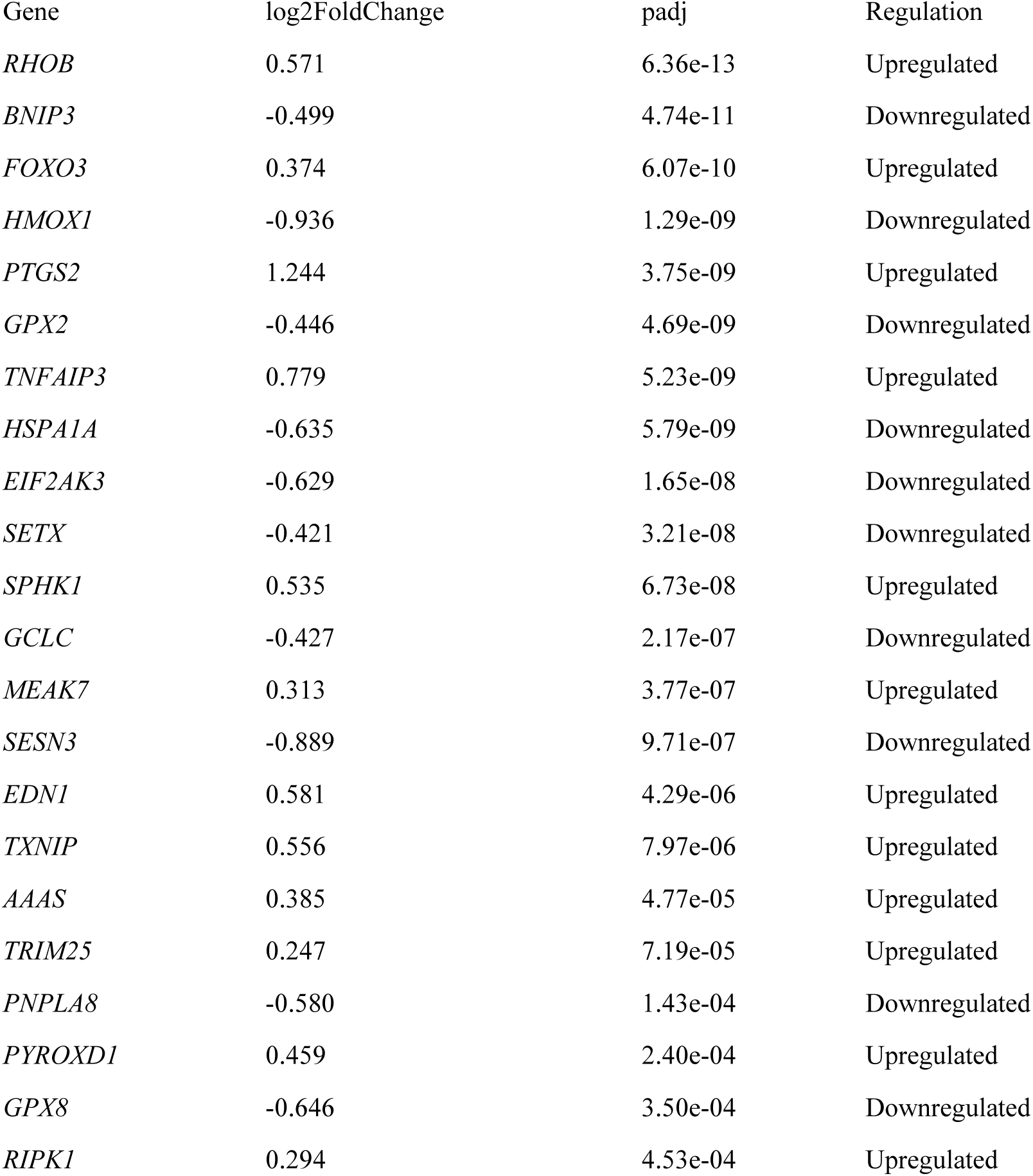
Differentially expressed genes associated with the gene ontology–biological process (GO-BP) term response to oxidative stress in KYSE-510 cells following exposure to 10 µM tenuazonic acid (TeA) for 6 h. The pathway was significantly enriched after TeA treatment compared to 0.1% DMSO control (p.adjust = 8.17 × 10⁻⁴). The table shows genes contributing to this enriched term, including log2FoldChange, adjusted p-values, and direction of regulation.

Grouped gene expression patterns indicate an oxidative stress-associated response characterized by the activation of stress- and inflammation-related signaling pathways, while several cytoprotective adaptation mechanisms are reduced. In particular, genes involved in redox-sensitive stress signaling, including *DUSP1, FOXO3, TXNIP*, and *RHOB,* as well as inflammatory and damage-associated signaling genes such as *PTGS2, TNFAIP3*, *RIPK1, EDN1*, and *SPHK1*, were upregulated (Bernardo et al., 2023; Liu et al., 2008; Martín-Vázquez et al., 2023; Prescott et al., 2021; Yoshihara et al., 2014). In contrast, several genes associated with antioxidant defense and ROS detoxification, including *HMOX1, GPX2, GPX8, GCLC*, and *SESN3*, were downregulated, suggesting an impaired compensatory antioxidant response (Ma, 2013; Pei et al., 2023). This was accompanied by a reduced expression of genes related to ER stress adaptation and proteostasis, including *XBP1, EIF2AK3, HSPA1A,* and *ATP2A2*, as well as mitochondrial stress adaptation genes such as *BNIP3* and *PNPLA8* (Gottlieb et Thomas 2017; Hetz et al. 2015, 2020). In parallel, *CYP1B1* was upregulated, indicating an involvement of xenobiotic- and stress-associated metabolic processes (Falero-Perez et al., 2018). Exposure to 1 µM TeA did not result in changes in the set of genes for *oxidative stress*.

#### Intrinsic apoptotic signaling in response to oxidative stress

The GO-BP term *intrinsic apoptotic signaling pathway in response to oxidative stress* was significantly enriched (p.adjust = 2.83 × 10⁻²) (Table 2). The corresponding enrichment plot (Figure 9) illustrates the overall enrichment of GO-BP term *intrinsic apoptotic signaling pathway in response to oxidative stress* (GO:0008631) within the ranked gene list.

**Figure 9:**
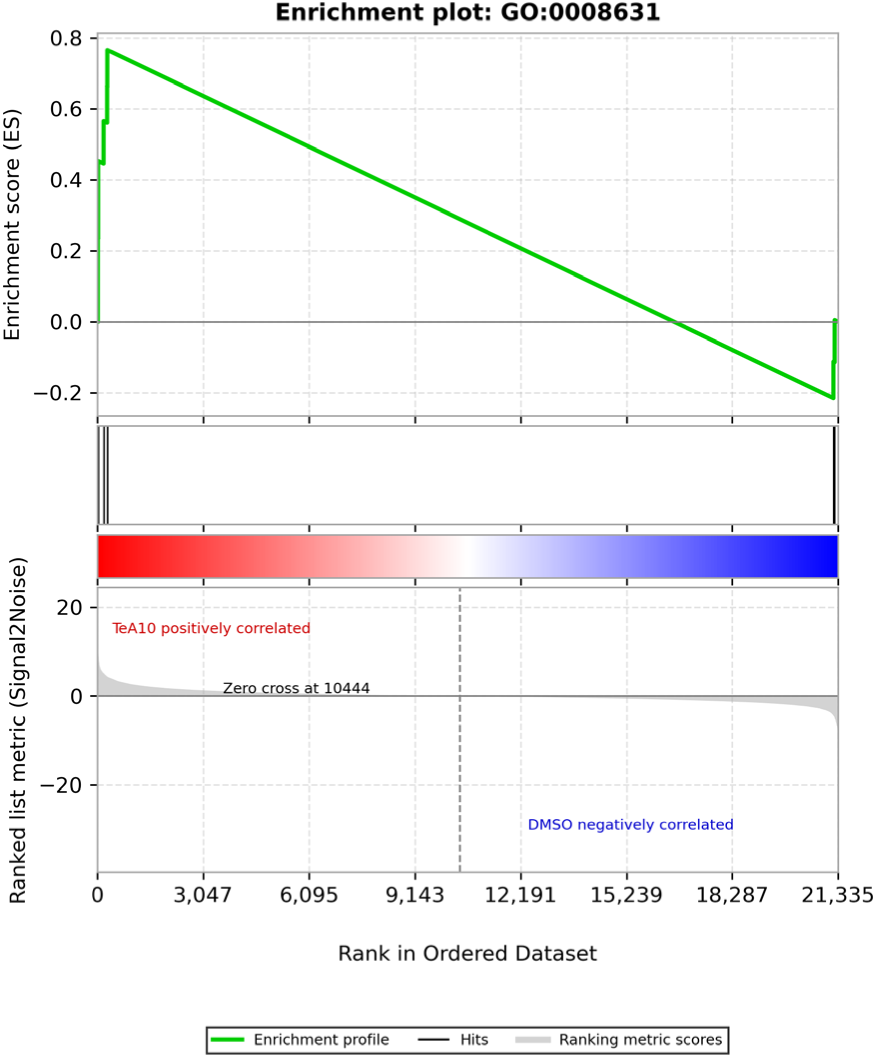
Gene set enrichment plot of the GO-BP term intrinsic apoptotic signaling pathway in response to oxidative stress (GO:0008631) in KYSE-510 cells after exposure to 10 µM tenuazonic acid (TeA) for 6 h compared with 0.1% DMSO control. Genes are ranked according to the signal-to-noise metric, with genes positively correlated with 10 µM TeA located on the left and genes negatively correlated with DMSO located on the right. Black vertical lines indicate the position of genes belonging to the GO:0008631 gene set within the ranked dataset, and the green line represents the running enrichment score. The plot illustrates the enrichment of this gene set toward the 10 µM TeA-correlated side of the ranked gene list.

**Table 2:** Differentially expressed genes associated with the GO-BP term intrinsic apoptotic signaling pathway in response to oxidative stress in KYSE-510 cells following exposure to 10 µM tenuazonic acid (TeA) for 6 h. The pathway was significantly enriched after TeA treatment compared to 0.1% DMSO control (p.adjust = 2.83 × 10⁻²). The table shows genes contributing to this enriched term, including log2FoldChange, adjusted p-values, and direction of regulation.

| Gene | $\log_2$FoldChange | padj | Regulation |
| --- | --- | --- | --- |
| *ZNF622* | 0.561 | 6.64e-22 | Upregulated |
| *CYP1B1* | 0.649 | 1.46e-18 | Upregulated |
| *MCL1* | 0.335 | 1.13e-05 | Upregulated |
| *P4HB* | -0.413 | 1.40e-05 | Downregulated |
| *PDK1* | -0.349 | 2.46e-04 | Downregulated |
| *NONO* | 0.204 | 2.73e-04 | Upregulated |
| *SFPQ* | 0.284 | 2.91e-04 | Upregulated |

The transcriptomic profile indicates a stress-associated response characterized by the upregulation of genes involved in anti-apoptotic/survival signaling, xenobiotic-associated metabolism, RNA processing and translational stress, including *MCL1, CYP1B1, NONO, SFPQ* and *ZNF622* (Falero-Perez et al., 2018; Hetz et al., 2020; Senichkin et al., 2020). In contrast, genes associated with ER redox homeostasis and metabolic adaptation, including *P4HB* and *PDK1*, were downregulated (F. He et al., 2020; Pei et al., 2023). Together, these data suggest an activation of stress-related survival and nuclear stress responses, accompanied by reduced ER redox and metabolic adaptation.

#### p53-associated signaling pathways

The GO-BP terms *signal transduction by p53 class mediator* (p.adjust = 1.53 × 10⁻³) was significantly enriched following TeA exposure (Table 3). The corresponding enrichment plot (Figure 10) illustrates the distribution of GO-BP terms *signal transduction by p53 class mediator* (GO:0072331)-associated genes within the ranked gene list and indicates enrichment toward the TeA10-correlated side.

**Figure 10:**
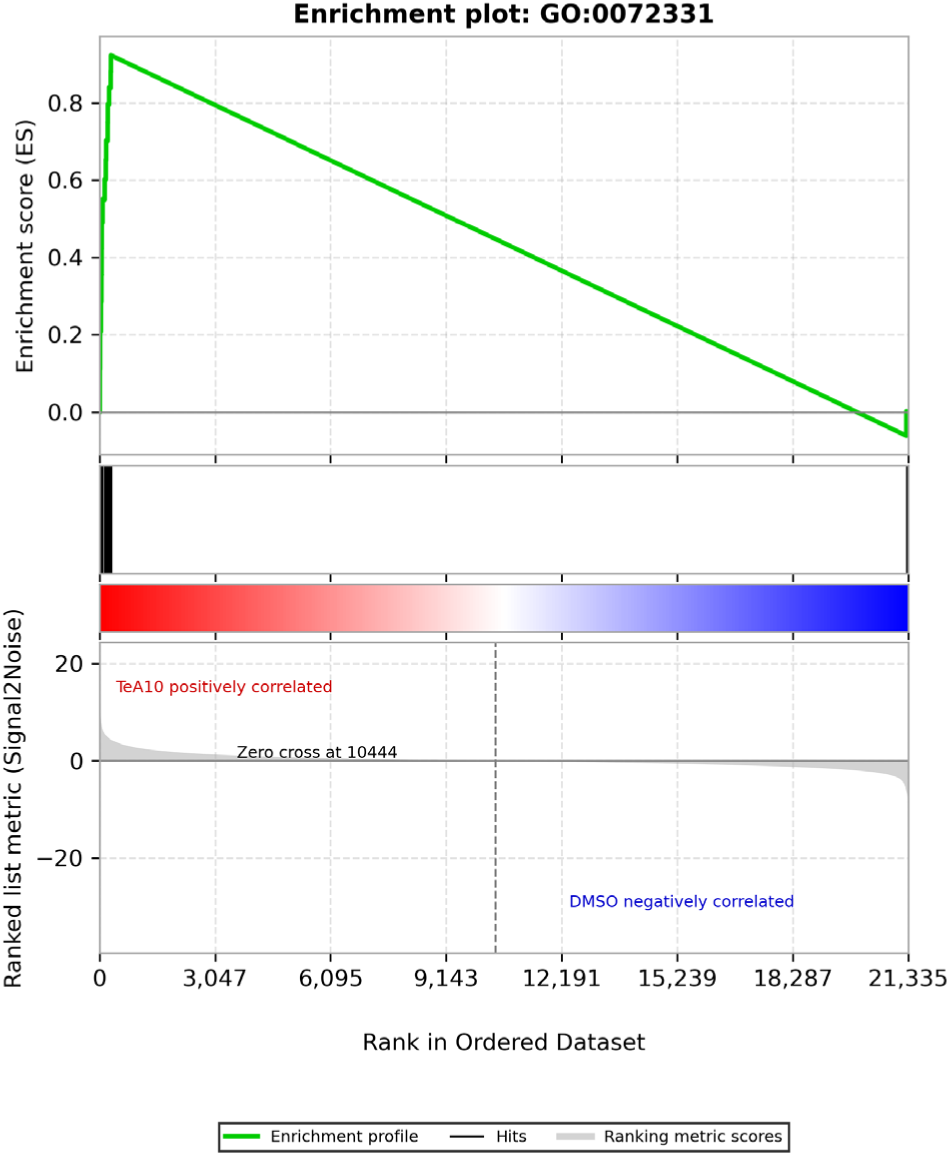
Gene set enrichment plot of the GO-BP term signal transduction by p53 class mediator (GO:0072331) in KYSE-510 cells after exposure to 10 µM tenuazonic acid (TeA) for 6 h compared with 0.1% DMSO control. Genes are ranked according to the signal-to-noise metric, with genes positively correlated with 10 µM TeA located on the left and genes negatively correlated with DMSO located on the right. Black vertical lines indicate the position of genes belonging to the GO:0072331 gene set within the ranked dataset, and the green line represents the running enrichment score. The plot illustrates enrichment of p53 class mediator-associated signal transduction genes toward the 10 µM TeA-correlated side of the ranked gene list.

**Table 3:** Differentially expressed genes associated with the GO-BP term signal transduction by p53 class mediator in KYSE-510 cells following exposure to 10 µM tenuazonic acid (TeA) for 6 h. The pathway was significantly enriched after TeA treatment compared to 0.1% DMSO control (p.adjust = 1.53 × 10⁻³). The table shows genes contributing to this enriched term, including log2FoldChange, adjusted p-values, and direction of regulation.

| Gene | log2FoldChange | padj | Regulation |
| --- | --- | --- | --- |
| *PPP1R15A* | 0.776 | 1.57e-28 | Upregulated |
| *NOP53* | 0.667 | 2.24e-20 | Upregulated |
| *DDX5* | 0.527 | 3.28e-13 | Upregulated |
| *SNAI2* | 0.743 | 1.01e-10 | Upregulated |
| *AEN* | 0.518 | 3.19e-10 | Upregulated |
| *FOXO3* | 0.374 | 6.07e-10 | Upregulated |
| *AARS1* | -0.348 | 7.30e-09 | Downregulated |
| *RRS1* | 0.676 | 2.60e-08 | Upregulated |
| *TP53BP2* | 0.337 | 2.37e-06 | Upregulated |
| *SNW1* | 0.413 | 6.72e-06 | Upregulated |
| *IFI16* | 0.410 | 9.12e-06 | Upregulated |
| *RRN3* | 0.301 | 4.16e-05 | Upregulated |
| *YJU2* | 0.497 | 4.34e-05 | Upregulated |
| *PHLDA3* | 0.395 | 1.30e-04 | Upregulated |
| *RPF2* | 0.486 | 3.83e-04 | Upregulated |
| *SOX4* | 0.298 | 4.08e-04 | Upregulated |

The observed gene expression pattern points towards a p53-related stress response with a strong involvement of nucleolar and transcriptional processes. In particular, the upregulation of *PPP1R15A, AEN, TP53BP2, PHLDA3* and *IFI16* suggests activation of stress signaling pathways linked to p53-mediated cellular responses (Fujiuchi et al., 2004; Kawase et al., 2009; O’Brien et al., 2014). At the same time, increased expression of *NOP53, RRS1, RPF2* and *RRN3* indicates alterations in ribosome biogenesis and nucleolar stress (Klinge et Woolford 2019), while *DDX5, SNW1* and *YJU2* point towards changes in RNA processing and transcriptional regulation (Li et Manley, 2009). The upregulation of *SNAI2* and *SOX4* further suggests transcriptional remodeling and cellular plasticity (Lourenço et Coffer 2017). In contrast, the downregulation of *AARS1* indicates that translation- and protein synthesis-related processes might be reduced (Gao et al., 2025).

## Discussion

TeA is a secondary fungal metabolite produced by *Alternaria* species and has been detected in several food commodities, including grains, oilseeds, tomatoes and fruits (Arcella et al. 2016; Asam et al. 2012; Asam et Rychlik 2013; Gotthardt et al. 2019). Available ADME data indicate rapid absorption, systemic distribution and predominant urinary elimination (Puntscher et al., 2019; Visintin et al., 2023).TeA has been associated with severe adverse effects and its toxicological relevance is further supported by inhibition of eukaryotic protein synthesis (Carrasco et Vazquez 1972, 1973; Farber et al. 1971; Kumari et Singh 2021; Lieberman et al. 1970; Louro et al. 2024; Shigeura et Gordon 1963; Younger 1970). Epidemiological data suggest an association between high dietary exposure to TeA-contaminated food and increased esophageal cancer incidence (Mulisa et al., 2025). This is supported by murine findings showing TeA-induced histopathological alterations in the esophageal mucosa, highlighting the relevance of investigating its toxicological effects in esophageal cell models (Yekeler et al., 2001).

In KYSE-510 cells, TeA induced a clear concentration-dependent reduction in cellular viability after 24 h of exposure. TeA concentrations up to 10 µM did not significantly affect cellular viability, whereas exposure to ≥20 µM resulted in a pronounced reduction in metabolic activity (Figure 2). Compared to other cell models, including Caco-2, HepG2, HepaRG and HeLa cells, KYSE-510 cells exhibited markedly higher sensitivity to TeA. Reported IC₅₀ values in these cell lines ranged from 109 to 308 µM (den Hollander et al., 2022; Hessel-Pras et al., 2019; Mahmoud et al., 2022), while cell viability in KYSE-510 cells decreased to 47% at 75 µM. Thus, the cytotoxicity data defined a concentration range in which early stress responses could be investigated below overt cytotoxicity.

Furthermore, exposure of KYSE-510 cells to TeA induced a pronounced but transient increase in intracellular ROS as determined by the DCF-DA assay. High TeA concentrations caused an immediate and significant ROS elevation after short-term exposure, whereas lower concentrations showed a delayed but significant increase (Figure 3). Oxidative stress is recognized as a central mechanism underlying the toxicity of several *Alternaria* mycotoxins. Alternariol and alternariol monomethyl ether have been shown to induce dose-dependent ROS formation in human cells, with growth inhibition correlating with ROS generation (Pahlke et al., 2016; Solhaug et al., 2012; Tiessen et al., 2017). In line with the observed ROS formation, *in vivo* data also indicate that TeA can induce oxidative stress. Kumari and Singh (2021) showed that sub-chronic TeA exposure in mice increased lipid peroxidation, reflected by elevated malondialdehyde levels, and impaired antioxidant defense, as indicated by reduced superoxide dismutase and catalase activities. Consistent with these findings, the present data suggest that TeA-induced cytotoxicity may be at least partly mediated by oxidative stress.

The rapid ROS formation observed after TeA exposure may be explained by its 3-acyl tetramic acid structure. This scaffold undergoes keto-enol tautomerism and can form a deprotonated enolate under physiological conditions. The resulting O,O-chelating motif enables the coordination of metal cations, including iron and copper (Poliseno et al. 2021; Zaghouani et Nay 2016). Therefore, TeA may disturb intracellular metal homeostasis rather than directly generating ROS itself. In particular, TeA-bound Fe/Cu complexes may increase the labile, redox-accessible metal pool and thereby facilitate endogenous redox reactions. This has been shown for N,N,N′,N′-tetrakis(2-pyridylmethyl)ethylenediamine (TPEN), which induces ROS formation and cell death through the formation of redox-active TPEN–copper complexes, indicating that metal chelation does not necessarily result in antioxidant effects but may also promote oxidative stress under certain conditions (Fatfat et al., 2014). In the presence of cellular reductants and H₂O₂, this could promote Fenton-like reactions or contribute to mitochondrial redox imbalance, resulting in rapid ROS accumulation (Abe et al., 2022; Georgieva et al., 2017). Thus, the early ROS response is most plausibly linked to TeA-mediated metal chelation and disturbance of Fe/Cu-dependent redox homeostasis, whereas inhibition of protein synthesis may represent an additional mechanism contributing to cellular stress. This mechanism might also be closely connected to mitochondrial dysfunction and impaired mitochondrial turnover, as mitochondrial homeostasis depends on continuous synthesis, import and replacement of mitochondrial proteins (Moehle et al., 2019; Richter et al., 2019). Since mitochondria have a high demand for protein renewal and quality control, disturbances in ribosomal activity or translational capacity may compromise mitochondrial adaptation under stress conditions. In line with this, deoxynivalenol exposure has been shown to induce a mitochondrial stress signature in human epidermal cells, suggesting that ribotoxic stress and mitochondrial impairment can be functionally connected (Del Favero et al., 2021). Furthermore, enhanced removal of damaged mitochondria by mitophagy reduced mitochondrial oxidative stress and increased tolerance to trichothecenes, supporting the relevance of mitochondrial quality control in the cellular response to ribosome-targeting mycotoxins (Bin-Umer et al., 2014).

To further assess whether TeA-induced oxidative stress resulted in DNA damage, the alkaline comet assay was performed with and without FPG treatment after 1 h of exposure. In the absence of FPG, TeA did not induce a clear increase in tail intensity, indicating that acute exposure did not result in pronounced DNA strand break formation under the investigated conditions. In contrast, FPG treatment significantly increased tail intensity at 5 and 7.5 µM TeA, reaching 13 ± 3 and 16 ± 3%, respectively, indicating the formation of FPG-sensitive DNA lesions (Figure 4). Since FPG mainly detects oxidized purines, including 8-oxoguanine-related lesions, these findings suggest that TeA primarily induces oxidative DNA base damage rather than direct strand breaks (Collins, 2014; Muruzabal et al., 2021). This is in line with the rapid ROS formation observed in the DCF-DA assay and supports oxidative stress as an early response to TeA exposure.

Given the involvement of ROS, mitochondrial integrity and function after TeA treatment were examined after 4 h of exposure. Low concentrations did not markedly affect mitochondrial morphology. In contrast, 7.5 µM TeA, still below overt cytotoxic levels, significantly reduced the total mitochondrial network area per cell. Morphological appearance of the organelles suggests a shift from an extended filamentous network to a more punctate and perinuclear distribution pattern. The reduction in mitochondrial network area in the absence of changes in overall fluorescence intensity and cell morphology indicates mitochondrial rearrangement rather than mitochondrial loss (Figure 5), interpretation which would be compatible with the outcome of the cytotoxicity assay (Figure 2). Such intracellular reorganization represents a typical cellular stress response, as oxidative stress is known to rapidly promote mitochondrial fission (Iqbal et Hood 2014). The transient ROS-inducing effect of TeA likely contributed to the observed alterations in mitochondrial morphology. The formation of a smaller, perinuclear mitochondrial network may reflect early apoptotic signaling or an adaptive stress response (Youle et Van Der Bliek 2012).

Mitochondrial respiration was assessed by measuring OCR using the Seahorse XF assay. TeA at 1–5 µM did not alter basal respiration or responses to mitochondrial inhibitors and uncouplers, indicating preserved mitochondrial function. At 7.5 µM TeA, basal OCR showed a slight reduction compared to the solvent control. ATP-linked respiration tended to be lower, as indicated by a more pronounced decrease in OCR following oligomycin treatment. Furthermore, the increase in OCR after CCCP-induced uncoupling appeared attenuated, suggesting a reduced maximal respiratory capacity and reserve capacity. Although these changes did not reach statistical significance (Figure 6), they align with the cytotoxicity assay results, where in the same concentration range a fluctuation started towards signal reduction (Figure 2A). Collectively, mitochondrial integrity and OCR data suggest that sub-cytotoxic TeA exposure affects mitochondrial homeostasis. The concurrent observation of altered mitochondrial morphology and a tendency towards reduced respiratory parameters indicates that mitochondria may represent early and sensitive targets of TeA. These findings are consistent with reports demonstrating disturbances in cellular energy metabolism following TeA exposure, including enhanced oxidative stress and ATP depletion in co-exposure models (Zhou et al., 2024). However, since the alterations in respiratory capacity did not reach statistical significance, these findings should be interpreted with caution.

The RNA-seq analysis performed after 6 h of exposure provided molecular support for the functional data and indicated a coordinated cellular stress response. The upregulation of redox-sensitive and inflammation-associated stress genes, including *DUSP1, TNFAIP3, PTGS2, TXNIP, FOXO3* and *CYP1B1*, is consistent with activation of MAPK-, NF-κB- and ROS-associated stress signaling (Table 1, Figure 8). In particular, the upregulation of *TXNIP* may further contribute to oxidative stress by limiting thioredoxin-dependent redox control, thereby supporting the interpretation of a disturbed intracellular redox state (Bernardo et al. 2023; Falero-Perez et al. 2018; Hoppstädter et Ammit 2019; Liu et al. 2008; Martín-Vázquez et al. 2023; Prescott et al. 2021; Vereecke et al. 2009; Yoshihara et al. 2014). Importantly, this stress response was not accompanied by a coordinated activation of antioxidant defense mechanisms. Instead, several cytoprotective genes, including *HMOX1, GCLC, GPX2, GPX8* and *SESN3*, were downregulated (Table 1). Since these genes are functionally linked to NRF2-regulated antioxidant defense, glutathione homeostasis and peroxide detoxification, their reduced expression suggests an insufficient compensatory antioxidant response despite clear evidence of ROS formation (F. He et al., 2020; Ma, 2013; Pei et al., 2023). This may also be linked to the known inhibitory effect of TeA on protein biosynthesis. Since antioxidant adaptation requires the de novo synthesis of cytoprotective enzymes, interference with translational processes could limit the cellular capacity to mount an adequate antioxidant defense response. In addition to oxidative stress-associated signaling, the transcriptomic data also indicated impaired ER stress adaptation and disturbed proteostasis. Downregulation of *XBP1, EIF2AK3* and *HSPA1A* suggests a reduced capacity to maintain proteostasis (Table 1), which is in line with the well-described ability of TeA to inhibit eukaryotic protein synthesis (Hetz et al., 2015, 2020). In parallel, enrichment of p53-associated GO terms and upregulation of *PPP1R15A, TP53BP2, AEN, NOP53, RRS1* and *RPF2* point towards activation of integrated stress response as well as nucleolar and ribosomal stress signaling (Table 3, Figure 10). This is mechanistically plausible, since disturbed protein synthesis and ribosome-associated stress can activate nucleolar stress pathways that converge on p53-associated cellular stress surveillance (Golomb et al., 2014; Hetz et al., 2020; Olausson et al., 2012). In addition, downregulation of *BNIP3* (Table 1) may indicate impaired mitophagy or mitochondrial quality control, further supporting the concept that TeA affects mitochondrial homeostasis (Gottlieb et Thomas, 2017).

Cell cycle progression was analyzed by flow cytometry following 24 h TeA exposure to assess sub-cytotoxic cellular effects. TeA induced an accumulation of KYSE-510 cells in the G₂/M phase. This effect was significant at 1-10 µM (Figure 7). To date, TeA has not been shown to exert direct genotoxic activity, with negative Ames tests and no reported DNA adduct formation (Louro et al., 2024; Schrader et al., 2006; Schwarz et al., 2012). Therefore, the G₂/M phase accumulation observed in the present study is most likely an indirect consequence of cellular stress rather than a direct interaction with DNA. One plausible mechanism involves oxidative DNA damage, as the pronounced ROS induction following TeA exposure may lead to oxidative lesions, thereby activating the G₂/M checkpoint to allow repair prior to mitotic entry. This interpretation is further supported by the FPG-comet assay, in which TeA induced FPG-sensitive DNA damage after 1 h of exposure. Since FPG-sensitive sites mainly reflect oxidized purine lesions, including 8-oxoG-related DNA damage, these data provide a direct downstream consequence of TeA-induced oxidative stress. In addition, inhibition of protein synthesis by TeA may impair the production of essential cell cycle regulators, such as cyclins and checkpoint proteins, resulting in delayed cell cycle progression (Lockhead et al., 2020). The concentration-dependent increase in the G₂/M population in the absence of a pronounced sub-G₁ fraction indicates a predominantly cytostatic effect at lower concentrations, characterized by growth arrest rather than immediate cell death. This is further supported by the absence of a clear induction of canonical p53 effector genes involved in terminal cell-cycle arrest or apoptosis, arguing against a full p53-driven apoptotic response at the investigated time point (Table 2). In line with this, enrichment of intrinsic apoptotic signaling was not accompanied by a clear pro-apoptotic transcriptional signature, and the upregulation of the anti-apoptotic regulator *MCL1* (Table 2) suggests that cells may still activate survival mechanisms to delay apoptotic commitment (Senichkin et al., 2020). Although TeA at higher concentrations induced G₂/M accumulation comparable to the positive control oxaliplatin, the relatively stable S-phase fraction suggests a less uniform and potentially reversible arrest. Thus, the induction of G₂/M phase arrest by TeA in KYSE-510 cells likely acts in concert with its cytotoxic effects by slowing cell cycle progression and limiting cellular proliferation. This checkpoint activation may further sensitize cells to apoptotic processes if stress-induced damage, particularly oxidative injury, is not adequately repaired(P. He et al., 2020; C. Zhang et al., 2015).

Taken together, the data delineate a coherent sequence of TeA-induced cellular events in KYSE-510 cells. TeA induced an early oxidative stress response within minutes, followed by FPG-sensitive oxidative DNA damage after 1 h and alterations in mitochondrial morphology after 4 h. The RNA-seq data obtained after 6 h further support the activation of redox-sensitive stress signaling, impaired antioxidant defense, disturbed proteostasis and p53-associated stress responses. Effects of early to intermediate kinetic were followed by G₂/M phase accumulation and reduced cellular viability after 24 h. Thus, TeA appears to induce a time-resolved stress response in KYSE-510 cells, in which early ROS formation and oxidative DNA damage align to mitochondrial alterations, transcriptomic stress adaptation, cell-cycle perturbation and eventually cytotoxicity at higher concentrations or prolonged exposure.

## Statements and Declaration

### Author contributions

Conceptualization: Dino Grgic, Doris Marko, Giorgia Del Favero; Methodology: Dino Grgic, Maximilian Jobst, Mariagiovanna Pais, Sonja Hager; Formal analysis and investigation: Dino Grgic, Maximilian Jobst, Mariagiovanna Pais, Nazmi Waesoh; Writing – original draft preparation: Dino Grgic; Writing – review and editing: all co-authors; Funding acquisition: Doris Marko; Resources: Doris Marko, Giorgia Del Favero; Supervision: Doris Marko, Giorgia Del Favero

## Acknowledgements

Graphical abstract was created in BioRender. Call, F. (2026) https://BioRender.com/0gtodqn

